# PREDICTING RISK OF INADEQUATE MICRONUTRIENT INTAKE WITH TRANSFERABLE MACHINE LEARNING MODELS

**DOI:** 10.1101/2025.04.08.647715

**Authors:** Vasiliki Voukelatou, Kevin Tang, Ilaria Lauzana, Manita Jangid, Giulia Martini, Saskia de Pee, Frances Knight, Duccio Piovani

## Abstract

Identifying populations at risk of inadequate micronutrient intake is useful for governments and humanitarian organizations in low– and middle-income countries to make informed and timely decisions on nutrition relevant policies and programmes. We propose a machine-learning methodological approach using secondary data on household dietary diversity, socioeconomic status, and climate indicators to predict the risk of inadequate micronutrient intake in Ethiopia and in Nigeria. We identify key predictive features common to both countries, and we demonstrate the model’s transferability from one country to another to predict risk of inadequate micronutrient intake in contexts where nationally representative primary data are unavailable.

## 1 Introduction

Micronutrient deficiencies (MND), a form of malnutrition, affect half of all children and two thirds of women worldwide and pose a significant threat to the health, development, and economic productivity of countries and their populations [1, 2]. Low quality, non-diverse diets provide inadequate quantities of essential micronutrients and are a direct cause of MND, particularly in low– and middle-income countries (LMICs) [3, 4]. In response, public policy and programmes aimed at reducing the burden of MND often seek to improve micronutrient intake through vitamin and mineral fortification^1^ of staple foods, supplementation, dietary diversification and other food systems interventions [6, 7, 8, 9].

Determining the need for and effectively designing policies and programmes to reduce the risk of inadequate micronutrient intake requires quantitative dietary intake data. These data are used by government agencies or other organizations, such as the United Nations World Food Programme (WFP), to assess the extent to which a population’s micronutrient intake meets recommended levels. Analysis of these data can also identify geographies or sub populations most vulnerable to inadequate intake, micronutrient intake gaps that need to be addressed and the contributions different interventions could have to improving intake [10].

Dietary intake can be measured directly using individual-level dietary recall surveys and is used to estimate the risk of inadequate micronutrient intake [11]. Unfortunately, due to the cost and complexity of measuring dietary intake, few LMICs collect or have access to current and nationally representative dietary intake data [10, 12]. Alternatively, dietary intake and the risk of inadequate micronutrient intake can be estimated using the food consumption module of many nationally representative Household Consumption and Expenditure Surveys (HCES), which are conducted every 3-5 years in many LMICs [13, 14]. This approach presents an alternative option for filling micronutrient intake data gaps in countries where recent and sufficiently detailed HCES data exist and there is capacity to conduct the analysis. However, this option cannot be applied where HCES data are inexistent, out of date, insufficiently detailed to allow dietary analysis or do not represent the current situation following a shock or as a result of prolonged fragility [14, 15]. We define these contexts as data-constrained, while contexts with HCES data availability as data-rich.

Due to the critical need for data on the risk of inadequate micronutrient intake to appropriately respond to the burden of MND, alternative approaches to fill evidence gaps are being explored, including the use of machine learning techniques. The application of machine learning for filling micronutrient intake data gaps remains in its early stages. For example, a recent study used socioeconomic and health data from the 2016 Ethiopian Demographic and Health Survey, and applied machine learning techniques to predict proxy indicators of micronutrient intake, including food group consumption and receipt of micronutrient supplements, among Ethiopian children aged 6–23 months. They were able to identify relevant predictors of proxy indicators of micronutrient intake, suggesting large routine surveys could be useful sources of information for predictive models aimed at filling micronutrient data gaps [16].

Machine learning has been applied more broadly in food security to model and forecast indicators such as the Food Consumption Score (FCS) and the reduced Coping Strategy Index (rCSI) [17, 18] and build a cross-country prediction model [19]. These studies have informed our methodological approach to predicting risk of inadequate micronutrient intake, particularly in data selection, feature engineering and modeling.

In this study, we develop a machine-learning methodological approach leveraging secondary data on household dietary diversity, in terms of household level consumption of foods from different food groups diversity (referred to as “food group diversity” in the rest of the paper), socioeconomic, and climate related data, to predict the risk of inadequate micronutrient intake in Ethiopia and Nigeria. We identify key predictive features common to both countries and assess the model’s transferability from one country to another, demonstrating the potential to predict risk of inadequate micronutrient intake in contexts where nationally representative primary data are unavailable.

## 2 Results

The results presented in this manuscript aim to address three research questions, providing evidence on the suitability of machine learning models for predicting the risk of inadequate micronutrient intake, for five micronutrients (i.e. zinc, folate, iron, vitamin A, vitamin B12) assessed both individually and in combination using an overall indicator:

- RQ1: Can the risk of inadequate micronutrient intake be predicted using features related to food group diversity, socioeconomic status, and climate, by applying machine-learning techniques?
- RQ2: Which features are most important in driving the predictions, and how does feature importance differ between countries, assuming RQ1 holds true?
- RQ3: Can a machine-learning model trained in one country predict risk of inadequate micronutrient intake in another country?

To address our research questions, we have developed a machine learning-based methodological approach to predict the risk of inadequate micronutrient intake at household level. This approach leverages features related to food-group diversity, socioeconomic status, and climate. We select Ethiopia and Nigeria, as they both experience high rates of malnutrition and fragile food systems which are susceptible to climate vulnerability, food insecurity, and conflict and they have similar data availability, meaning matching features can be generated from the HCES data collected. An overview of the machine-learning methodological approach is presented in Figure 1.

**Figure 1:**
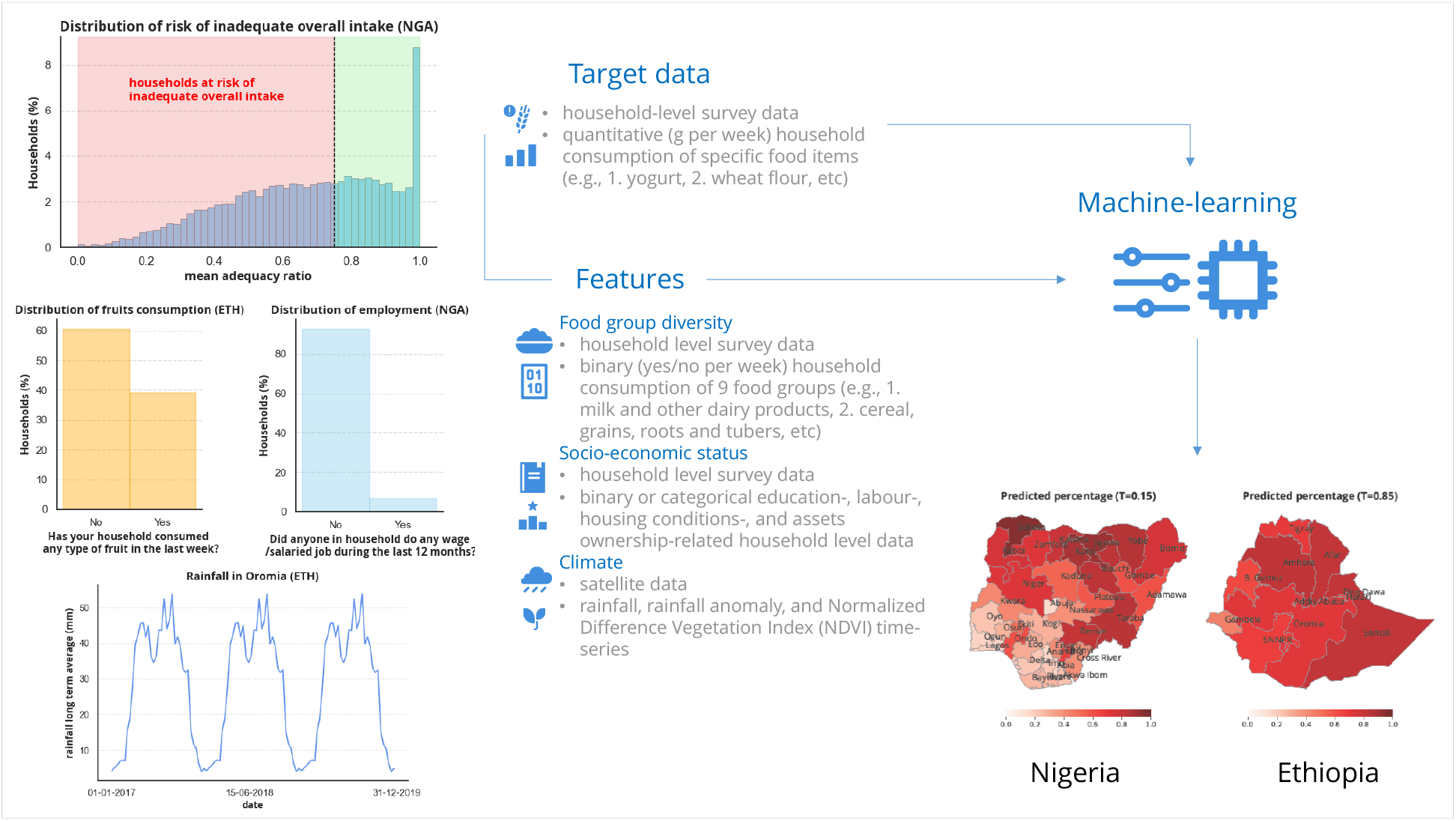
Overview of the machine-learning methodological approach: We extract quantitative food consumption data from household level surveys to estimate the target variables. From the same surveys, we derive features related to food group diversity and socioeconomic status. We also extract climate features from the WFP’s Seasonal Explorer platform. We then train machine-learning models using these variables to predict the risk of inadequate micronutrient intake.

### 2.1 Predicting risk of inadequate micronutrient intake with machine-learning

To address the first research question (RQ1), we trained an XGBoost binary classification model for the five micronutrients and the overall risk, separately for Ethiopia and Nigeria, meaning that the models are trained and tested within each country individually. Model performance is evaluated using the ROC-AUC score, which measures the model’s ability to discriminate between classes across a range of sensitivity and specificity thresholds that may vary depending on the application context. Table 1 presents the performances of the models for all target micronutrients in both countries, where the number in parenthesis indicates the normalized difference, i.e. percentage improvement in performance compared to a dummy model used as a benchmark, and ETH and NGA represent the ISO3 codes for Ethiopia and Nigeria, respectively.

**Table 1:** ROC-AUC scores, and the corresponding normalized difference (in parenthesis) of the XGBoost scores to the dummy models’ corresponding scores.

|  |  | zinc | folate | iron | vitamin A | vitamin B12 | overall risk |
| --- | --- | --- | --- | --- | --- | --- | --- |
| XGBoost |  |  |  |  |  |  |  |
| ETH | ROC-AUC | 0.76 | 0.77 | 0.76 | 0.79 | 0.87 | 0.82 |
| | ($\Delta$ ROC-AUC) | (0.51) | (0.54) | (0.53) | (0.57) | (0.73) | (0.63) |
| XGBoost |  |  |  |  |  |  |  |
| NGA | ROC-AUC | 0.71 | 0.72 | 0.73 | 0.75 | 0.77 | 0.77 |
| | ($\Delta$ ROC-AUC) | (0.43) | (0.44) | (0.45) | (0.51) | (0.54) | (0.54) |

Overall, the results indicate that the models in both countries outperform their respective dummy models by on average 0.59 and 0.49 in Ethiopia and Nigeria, respectively. We observe that model performances are consistent across models, with minor differences likely due to country-specific factors (e.g., local dietary patterns, and food availability) and the distribution of households at risk of micronutrient inadequacy (see Appendix E).

Next, we address the second research question (RQ2) to identify which variables used in the models contribute to a higher risk of inadequate micronutrient intake and which mitigate it, and how does this vary across the two countries.

For this purpose we use the SHAP methodology [20, 21], that allows to explain machine learning models by showing how much each feature contributes to a prediction. In particular, SHAP values represent the contribution of each feature to the model’s predicted probability of inadequacy or adequacy, with predictions classified based on the threshold *T* = 0.5. Figure 2 presents the ten most important features for the models trained to predict the risk of inadequate overall intake in Ethiopia (top) and Nigeria (bottom), ordered by their effect on the predictions. A positive SHAP value indicates that the feature increases the predicted outcome, while a negative SHAP value implies that the feature decreases it. All data points on the plot represent a single household observation, while the color shows a higher or a lower feature value (pink or blue, respectively). Thus, for example, from the figure we observe that reporting consumption of meat, fish or eggs (pink dots) results in negative SHAP value in both countries, meaning that it reduces the risk of inadequate overall intake, and vice versa for households that did not report consuming it. In general, comparing the feature importance for both countries we observe similar patterns, despite some ranking differences that are mostly observed towards the bottom of the list. For both country models, food group diversity-, climate– and education-related features rank among the most important features. Moreover, with few exceptions, the directionality of their impact is largely consistent between the most important features of the models (see Figure 2, and Appendix A for the SHAP values of each individual micronutrient model).

**Figure 2:**
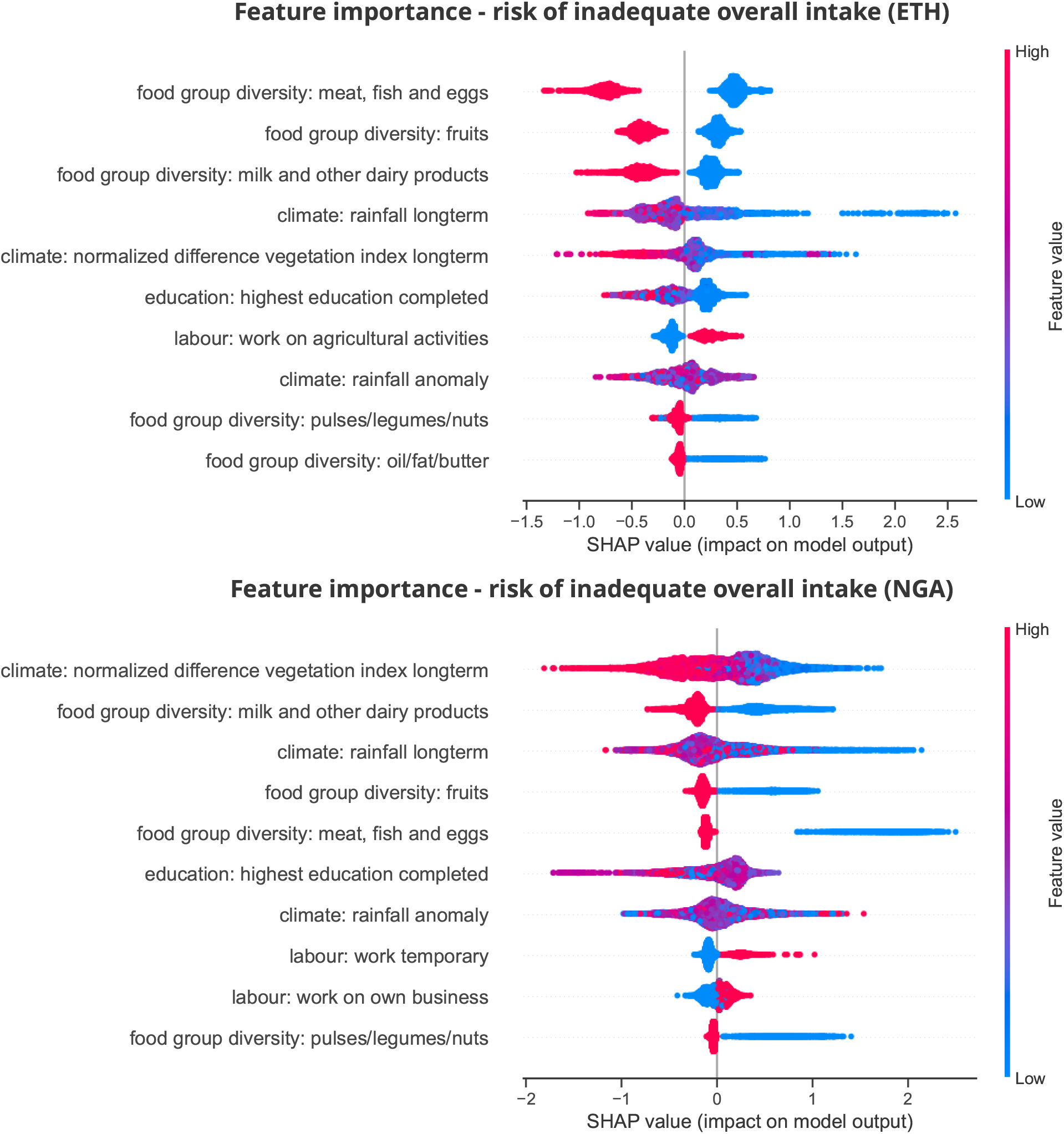
Feature importance for the risk of inadequate overall intake models, for Ethiopia (top) and Nigeria (bottom) country models. Feature importance remains consistent among the two models, despite some ranking differences, that are mostly observed towards the bottom of the list. The directionality impact is largely consistent between the most important features of the two models.

Overall, the robustness of the predictive models and the consistency in feature importance across both countries suggest that predictions are driven by similar underlying factors, despite potential contextual differences. This indicates that models developed in one country may be transferable to others, enhancing their applicability across different settings.

### 2.2 Cross-country models

In this section, we study whether a model trained in one country can predict risk of inadequate micronutrient intake in another country (cross-country model), addressing the third research question (RQ3). In this case, we treat Ethiopia and Nigeria, two data-rich countries, as data-constrained each time we train a model in one country and predict on the other. Similarly to the models trained at a country level, we build classification models, this time using 100% Ethiopian data for training and 100% Nigerian data for testing, and vice versa.

Table 2 reports the ROC-AUC scores of both cross-country models across all target micronutrients, along with the normalized differences (in parentheses) between XGBoost and dummy model scores. While the performance improvement is smaller compared to the models built within a country, the results indicate that all cross-country models still outperform their respective dummy models, with an average improvement of 0.21 and 0.35 for the model trained in Ethiopia and in Nigeria, respectively. This suggests that the models could serve as useful tools for preliminary risk assessments in data-constrained contexts. Similarly to Section 2.1, ROC-AUC scores show differences across models trained for different micronutrients, and countries. This might be attributed to country-specific factors and the distribution of households at risk of micronutrient inadequacy (see Appendix E).

**Table 2:** ROC-AUC score, and the corresponding normalized difference (in parenthesis) of the XGBoost scores to the dummy models’ corresponding scores.

|  |  | zinc | folate | iron | vitamin A | vitamin B12 | overall risk |
| --- | --- | --- | --- | --- | --- | --- | --- |
| XGBoost |  |  |  |  |  |  |  |
| ETH to NGA | ROC-AUC<br>($\Delta$ ROC-AUC) | 0.58<br>(0.16) | 0.58<br>(0.16) | 0.57<br>(0.14) | 0.61<br>(0.22) | 0.64<br>(0.28) | 0.64<br>(0.28) |
| XGBoost |  |  |  |  |  |  |  |
| NGA to ETH | ROC-AUC<br>($\Delta$ ROC-AUC) | 0.63<br>(0.26) | 0.61<br>(0.22) | 0.59<br>(0.18) | 0.68<br>(0.36) | 0.83<br>(0.66) | 0.71<br>(0.42) |

### 2.3 Application in data-constrained contexts

When applying this methodology in data-constrained contexts, where no ground-truth data is available, to validate predictions, alternative approaches are necessary. In such contexts, sanity checks can be employed to verify whether the predictions align with known patterns or distributions. To replicate this scenario, rather than simply relying on performance metrics, we assess cross-country predictions by reproducing and comparing two distributions observed in the actual data from the models’ outputs: the percentages of risk of micronutrient inadequacy and adequacy stratified by wealth quintile and by rural and urban divide (see more details in J). The first observed pattern, is that households in higher wealth quintiles tend to have a higher percentage of micronutrient adequacy compared to those in lower wealth quintiles [22], while the second observed pattern is that urban households generally exhibit greater micronutrient adequacy than rural households [23]. However, there is often substantial heterogeneity among urban populations, and country-specific contextual factors should be considered.

To reproduce the distributions observed in the actual data, the cross-country models presented in Section 2.2 need to be calibrated by adjusting the probability threshold that determines whether a household is classified as at risk of inadequate overall intake. A threshold of *T* = 0.5 means that a household is considered at risk, if the model assigns it a probability greater than 0.5. This means that higher thresholds make the model more conservative, while lower thresholds increase its sensitivity. Figure 3 presents the results of the model trained in Ethiopia and applied in Nigeria (3a-3c), and of the model trained in Nigeria and applied in Ethiopia (3d-3f), when modeling for the overall risk. Figures 3a and 3d show the ROC-AUC curve, reflecting the balance of true positive and false positive rates, across all possible probability thresholds. Figures 3b and 3c, and Figures 3e and 3f present the percentages of micronutrient inadequacy and adequacy when stratifying by wealth quintile and rural-urban area, for the model applied in Nigeria and the model applied in Ethiopia, respectively. Distributions of the predicted outputs are shown at three probability thresholds (*T* = 0.15, *T* = 0.5, and *T* = 0.85), compared to the actual data. Notably, at *T* = 0.15 for the model applied in Nigeria, and at *T* = 0.85 for the model applied in Ethiopia, the distributions of the predicted outputs closely align with the actual data: in the highest wealth quintile, micronutrient adequacy is higher than inadequacy, and higher in urban areas compared to rural areas. This suggests the need for a more reactive model when training in Ethiopia and applying in Nigeria, and a more conservative model when training in Nigeria and applying in Ethiopia. This may be due to differences in feature distributions between the two countries, pushing the model trained in Nigeria to overestimate the risk, while the model trained in Ethiopia underestimates it (see details in Appendix F).

**Figure 3:**
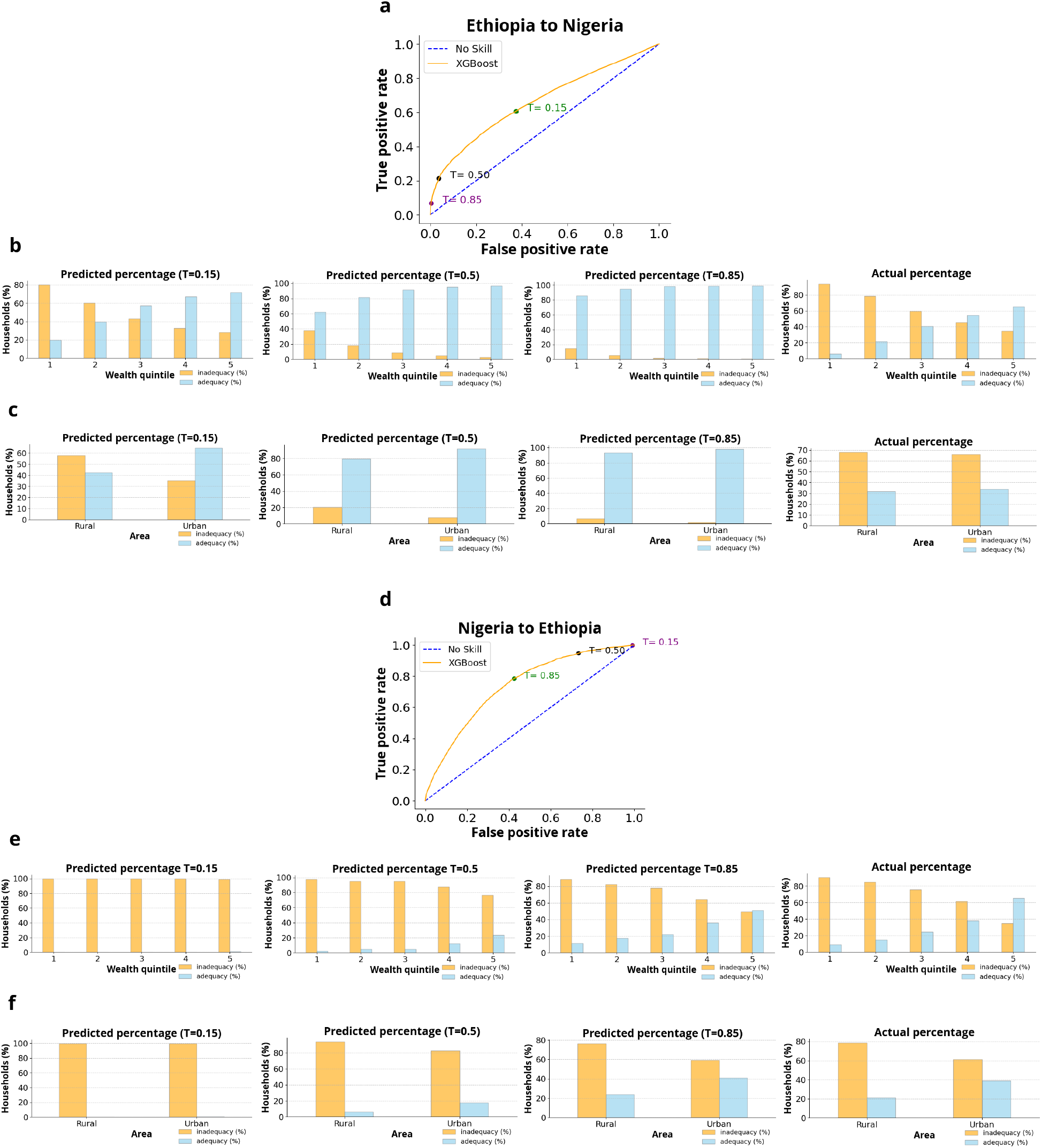
The Figure presents the results of the model trained in Ethiopia and applied in Nigeria (3a-3c), and of the model trained in Nigeria and applied in Ethiopia (3d-3f), when modeling for the overall risk. Figures 3a and 3d show the ROC-AUC curve, reflecting the balance of true positive and false positive rates, across all possible probability thresholds. Figures 3b and 3c, and Figures 3e and 3f present the percentages of micronutrient inadequacy and adequacy when stratifying by wealth quintile and rural-urban area, for the model applied in Nigeria and the model applied in Ethiopia, respectively. Distributions of the predicted outputs are shown at three probability thresholds (*T* = 0.15, *T* = 0.5, and *T* = 0.85), compared to the actual data. Notably, at *T* = 0.15 for the model applied in Nigeria, and at *T* = 0.85 for the model applied in Ethiopia, the distributions of the predicted outputs closely aligns with the actual data.

Figure 4 presents the actual and predicted risk of inadequate overall intake by first-level administrative unit resolution (see Appendix J). The optimal probability thresholds of both cross-country models yield predicted patterns that closely align with the actual patterns. Although the predicted percentages are not identical to the actual percentages, the results effectively identify areas of higher and lower risk (see Appendix G). For instance, in Figure 4a, the predicted percentages indicate a higher overall risk in northern Nigeria compared to southern Nigeria, consistent with the actual percentages. Thus, we observe that using sanity checks to determine the optimal threshold does not only produce a more interpretable output but also improves the spatial accuracy of the predictions.

**Figure 4:**
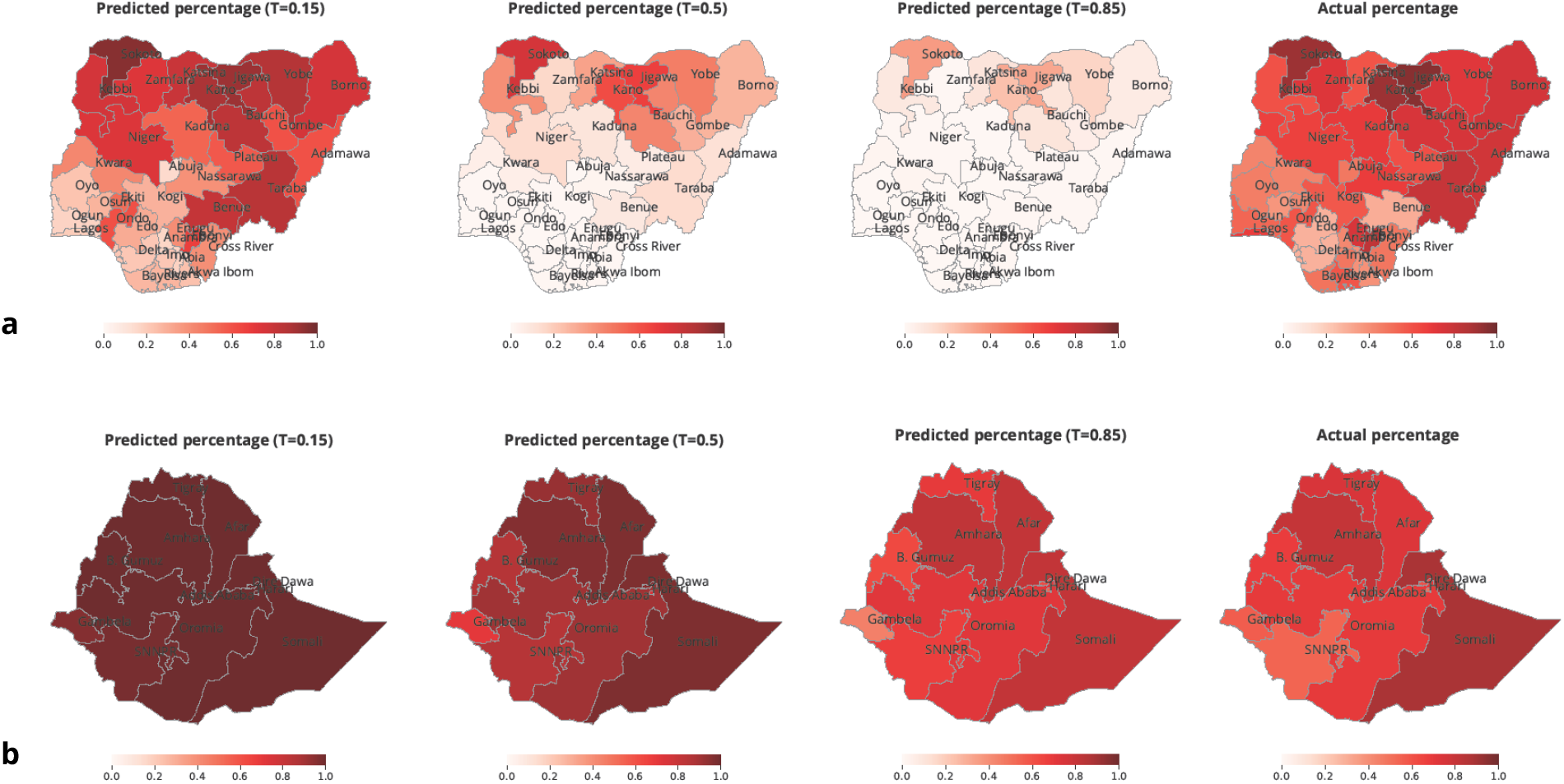
Actual and predicted percentages of risk of inadequate overall intake for the cross-country models applied in Nigeria (4a) and Ethiopia (4b) respectively, at first-level administrative unit resolution. We notice that for the optimal probability thresholds, i.e., at *T* = 0.15 for the model applied in Nigeria, and at *T* = 0.85 for the model applied in Ethiopia, the predicted percentages are very similar to the actual percentages.

## 3 Discussion

In this study, we demonstrate that using food-group diversity, socioeconomic, and climate data, and applying machine-learning techniques enables the prediction of risk of inadequate micronutrient intake. We show that drivers of inadequate micronutrient intake are similar, both in importance and directionality, in two different countries: in Nigeria, a West African country, and in Ethiopia, a Horn African country. These drivers are interpretable and align with established patterns identified in prior research [24, 25]. We also demonstrate that a model trained in one country can be potentially applied to predict inadequate micronutrient intake in another country. We simulate a scenario to determine the optimal probability threshold, achieving the right balance between true positive and false positive rate, to generate predictions of risk of inadequate micronutrient intake, which when stratified by wealth quintile and rural-urban area, exhibit the expected patterns.

Despite the encouraging results, several limitations apply. Although we have demonstrated that the feature importance is very similar across two countries, a larger training dataset including more countries, could further ensure that the models capture patterns that are most commonly observed across multiple countries. This would enable the assessment of the possibility of building a global model, which could be used to predict micronutrient inadequacy in other LMICs, or building regional models, which capture patterns specific to certain geographical contexts. However, it is important to note that before building the machine-learning models, target variables needed for the training must be estimated, a process that requires collaboration with relevant stakeholders in each country and access to the most recent national survey data.

Additionally, future research should focus on enhancing model performance by exploring more advanced algorithms and expanding the range of input features. This could include integrating higher-resolution climate data which capture variation below the second-level administrative unit resolution, as well as incorporating features related to conflict, market accessibility, and food prices. Furthermore, generative AI models could be leveraged to extract relevant insights from localized unstructured data sources, such as news reports and media mentions [26]. Moreover, rather than relying only on SHAP values to assess feature importance, a more rigorous causal analysis could help identify the direct effects of individual features on the risk of micronutrient inadequacy, offering clearer insights into cause-and-effect relationships between features and the target variable.

Future work also entails real-world application of the suggested methodological approach in data constrained contexts where HCES data are not available, and thus target data cannot be estimated. In such contexts, other available survey data can be used to generate food group diversity, and socioeconomic features, while climate features can be still extracted from the WFP’s Seasonal Explorer platform [27]. Table 6 in Appendix L provides the data sources, and availability for both data-rich and data-constrained contexts for all variables. It is important to emphasize that the application of the model in a real, data-constrained context should be treated carefully: feature distributions should be considered, and sanity checks should be guided by domain expertise. For example, while higher wealth is generally associated with improved micronutrient adequacy, urban settings can sometimes present mixed effects due to varied dietary patterns, including increased access to ultra-processed foods and a higher consumption of foods away from home. Therefore, while the expected trends generally hold, additional country-specific contextual factors should be considered in interpreting results. Predictions generated should also be interpreted with caution, and to ensure the validity of the proposed models, it is essential to triangulate the results with other available nutrition data sources. The predicted percentages of risk of micronutrient inadequacy should serve as insights, to better understand the context and make a case for more in-depth assessments as a next step.

After thorough consideration of the challenges and limitations mentioned above, the presented methodological machine learning approach could contribute as a detection tool to foster data-driven nutrition decision making when data are not available to estimate the risk of inadequate micronutrient intake.

## 4 Data and methods

This section outlines the variables, both targets and features, and the methods used in the proposed machine learning approach to predict the risk of inadequate micronutrient intake. Details on target variables and predictive features are provided in Sections 4.1 and 4.2, respectively. The modeling approach and the SHAP method are presented in Sections 4.3 and 4.4, respectively. An overview of the machine-learning methodological approach is presented in Figure 1.

### 4.1 Target data

We focus on five micronutrients of particular public health importance: zinc, folate, iron, vitamin A, and vitamin B12 [2]. Thus, we generate six target variables, one for each of the five micronutrients mentioned above and one for the risk of inadequate overall intake. The overall risk of inadequacy is assessed by calculating a Mean Adequacy Ratio (MAR) as an indicator of general dietary quality that averages the adequacy (% of estimated average requirement) across the five focus micronutrients [28]. The criteria for the micronutrient selection and the MAR calculation are detailed in Appendix B.

To generate the target variables, we use data from Household Consumption and Expenditure Surveys (HCES) collected as part of the Living Standards Measurement Study (LSMS) in both Ethiopia and Nigeria. HCES are a family of nationally representative cross-sectional surveys which aim to collect microdata that describe consumer behaviour and social welfare within a country. For Ethiopia, we use the fourth wave of the Ethiopia Socioeconomic Survey, a nationally representative survey which collected information from 6770 households between May to September 2019 [29]. For Nigeria, we use the Nigerian Living Standards Survey, a nationally representative survey which collected information from 22,587 households between September 2018 to September 2019 [30].

Specifically, we use the food consumption modules from each survey to estimate the risk of inadequate intake for each of the five focus micronutrients. First, we generate estimates of apparent micronutrient intake for each micronutrient, at a household level, maintaining consistency with methods established in prior research [31]. Both Ethiopian and Nigerian HCES collect food consumption data using a fixed list of commonly consumed food items (72 in Ethiopia, 113 in Nigeria) consistent with a standardised approach outlined in published guidance [32, 33]. To generate these estimates of apparent micronutrient intake, food items are matched to corresponding food composition values from locally relevant databases [34], micronutrient contents are summed across food items, and then proportionally redistributed to household members using the adult female equivalent (AFE) approach [35].

To classify the household level estimates (continuous values) into binary indicators of risk of inadequate micronutrient intake and inadequate overall intake, we compare each household’s estimated apparent intake per AFE to the corresponding Harmonized Average Requirement (H-AR) for an 18–29-year-old, non-pregnant, non-breastfeeding female [36]. The H-AR serves as the scientific threshold for determining whether a household’s intake is classified as adequate or inadequate for each individual micronutrient and for the MAR. For iron and zinc specifically, H-AR are set assuming low bioavailability, ensuring a conservative estimate of inadequacy. A household is classified as at risk of inadequate intake for a given micronutrient if its apparent intake falls below the respective H-AR, following the cut-point approach [37]. To compute MAR we first calculate the Nutrient Adequacy Ratio (NAR) for each of the five micronutrients. This is the ratio of the household’s apparent intake to the respective H-AR, with values truncated at 1 [28]. The MAR is then computed as the mean of the five NARs, yielding a continuous score ranging from 0 to 1. To classify households at risk of inadequate overall intake, we use a threshold of MAR *<* 0.75, meaning households which meet less than 75% of the Estimated Average Requirement (EAR) are considered at risk. This threshold is the recommended nutritional objective for the five included micronutrients used in nutrition programme planning in LMICs for increasing micronutrient intakes through interventions including food fortification, according to global guidance [38]. The distributions of the generated targets, for both Ethiopia and Nigeria, can be found in Appendix E.

### 4.2 Feature selection and engineering

Prior research has linked micronutrient inadequacy and its determinants to diversity of food groups consumed, socioeconomic [24] and climate indicators [25]. Thus, we select a set of features related to the indicators mentioned above to predict risk of micronutrient inadequacy. These features can be extracted from secondary data sources available in many countries. The initial set of considered features is composed of 25 features related to the aforementioned drivers. Considering that it is important to provide interpretable machine-learning insights to the relevant stakeholders, feature selection and engineering were carefully studied and guided by experts. The feature engineering conducted is not expected to remove multi-collinearity, but the XGBoost model used for modeling is robust to multi-collinearity. More details on feature engineering are provided below:

#### Food group diversity features

To generate food group diversity-related features, we use data from a different module of the same survey used to generate the targets (see Section 4.1), specifically the HCES data collected as part of the LSMS 2018/19 for Ethiopia and Nigeria [29, 30]. In the survey module used, intended to construct the Food Consumption Score [39], respondents report whether household members consumed foods from specific food groups, such as “cereal, grains, roots and tubers, etc”, during the previous week, using a binary yes/no format. In contrast, the data used to generate the targets involve quantitative reporting of household level consumption of specific food items, such as “wheat flour” measured in grams per week. These methodological differences are further detailed in Figure 1. Food group diversity features represent the household dietary diversity score as a proxy for dietary quality. We generate nine binary food group diversity features from each survey, at a household level. For example, one binary feature indicates whether a household consumed fruits in the past week. We align the food groups of the corresponding survey based on the Food Consumption Groups (FCGs), as defined by the United Nations WFP [39]. Table 3 in Appendix C shows the nine FCGs, and examples of food groups derived from each LSMS survey, which fall under the FCGs.

**Table 3:** United Nations World Food Programme Food Consumption Groups for the dietary diversity score, and examples of food groups from the Living Standards Measurement Study 2018/19 that fall under the WFP food groups, both for Ethiopia (ETH) and Nigeria (NGA).

| WFP FCGs [39] | LSMS (ETH)<br>food groups [29] | LSMS (NGA)<br>food groups [30] |
| --- | --- | --- |
| cereal, grains,<br>roots and tubers | e.g, roots and tubers, etc | e.g, grains and flours, etc |
| pulses/legumes/nuts | pulses, nuts and seeds | pulses, nuts and seeds |
| milk and<br>other dairy products | milk/yogurt/<br>cheese/other dairy | milk/milk products |
| meat, fish and eggs | e.g, poultry, eggs,<br>fish and seafood, red meat, etc | meat, fish and<br>animal products |
| vegetables and leaves<br>(including relish and leaves) | vegetables | vegetables |
| fruits | fruits: (mango; banana;<br>orange; pineapple; papaya;<br>avocado; other fruit) | fruits |
| oil/fat/butter | oils/fats/butter | oils and fats |
| sugar, or sweet | sweets (sugar or sugar<br>products (honey, jam) | sugar/sugar products/honey |
| condiments/spices | spices, condiments, beverages<br>(spices (pepper, salt), condiments,<br>coffee, tea, alcoholic beverages) | spices/condiments |

#### Socioeconomic features

To generate the socioeconomic-related features we use data extracted from the same survey used to generate the targets (see Section 4.1) and generate the food group diversity-related features, i.e., the HCES data collected as part of the LSMS 2018/19 [29, 30] for Ethiopia and Nigeria, respectively. We generate fourteen education-, labour-, housing conditions-, and assets ownership-related features. To ensure consistency of the features generated across the two different country data sets, we extract data collected from the same or similar questions. All socioeconomic features are generated at the household level. Housing conditions-, and assets ownership-related features are already collected at household level. Education– and labour-related data are collected at an individual household-member level, thus to generate the corresponding features at the household level, individual-level data are aggregated as following: labour-related data by taking the maximum value across all individuals in the household, while education-related data by calculating the median value of all individuals in the households. This distinction is made to enhance model interpretability and align with prior studies. Null values are imputed with 0 for all features, except for the education feature, for which the k-means algorithm is applied to maintain interpretability [40]. Examples of features are: an ordinal education-related feature varying from 0 to 6 representing the median value of the highest educational level completed in a household, a binary labour-related feature representing the maximum value over all individuals in the household on whether they were employed in last 12 months, etc. The detailed list of the features generated, and the corresponding survey questions can be found in Appendix D.

#### Climate features

To generate the climate-related features we extract data from WFP’s Seasonal Explorer platform [27], originally derived from satellite data. It provides open rainfall and Normalized Difference Vegetation Index (NDVI) time series for a near-global set of administrative units, computed, from the Climate Hazards Group InfraRed Precipitation with Station data (CHIRPS) [41] and the Moderate Resolution Imaging Spectroradiometer (MODIS) [42] data. We generate three climate-related features: (1) rainfall, by calculating the median of the 2017 to 2019 historical averages of rainfall, (2) NDVI, by calculating the median of the 2017 to 2019 historical averages of NDVI, and (3) rainfall anomaly, by calculating the median of 2017 to 2019 rainfall anomaly values with respect to historical averages. Since spatial resolution varies between HCES and climate data, climate features are built at second-level administrative unit resolution, leading to all households in the same administrative units being assigned the same value.

### 4.3 Modeling approach

The proposed models predict the probability of a household to be at risk of inadequacy of a single micronutrient, i.e., zinc, folate, iron, vitamin A, vitamin B12, or overall risk, with a separate model developed for each target (see 4.1 for more details on the targets). In total, we build twenty-four binary classification models. We develop twelve models trained within a country (six for Ethiopia and six for Nigeria). Also, we develop twelve cross-country models (six trained in Ethiopia and tested in Nigeria, and six trained in Nigeria and tested in Ethiopia). Each model corresponds to one of the six targets: five single micronutrients and the overall risk. The predictive features given as input to all models are the same: (1) food group diversity-, (2) socioeconomic-, and climate-related features (see 4.2 for more details on feature selection and engineering).

We build the models with the following methodological approach. We use the XGBoost algorithm given its high performance, and its capacity to handle complex and nonlinear relationships (e.g., [43], see Appendix H). When building the models at a country level, we use 80% of the data for training and the remaining data for testing, and we fine-tune the hyperparameters via 5-fold cross-validation. The tuned hyperparameters and the explored values are listed in Appendix I. Given that all targets are imbalanced we use an undersampling technique for all models. The undersampling technique randomly removes from the training data records that belong to the majority class in order to balance the class distribution, reducing the skew to an even 1:1 class distribution. For cross-country modeling, similarly to modeling within one country, we build classification models, this time by using 100% of Ethiopian data for training and 100% of Nigerian data for testing, and vice versa. Once the models are trained, we use the test set to evaluate our models.

To evaluate the performance of the models we use the area under the ROC-AUC curve, i.e. the ROC-AUC score, which assesses the models’ discriminative power between a positive or negative class. It shows the trade-off between the true positive rate (i.e., recall or sensitivity) and the false positive rate (i.e., specificity) for different probability thresholds [44]. The probability threshold (*T*) determines the cutoff point for classifying an observation as positive (household at risk of inadequacy) or negative (household not at risk of inadequacy). By default, classifiers set this threshold at *T* = 0.5, meaning predictions with a probability *T >* 0.5 are assigned to the positive class, while those at *T <*= 0.5 are assigned to the negative class. Lastly, to control the robustness of the machine-learning results, the data are resampled five times with randomization of the resampling procedure.

### 4.4 SHAP method

Understanding the models’ predictions is important for trust, accountability, and debugging. To understand predictions from tree-based machine learning models, like XGBoost, feature importance values are typically attributed to each feature. Yet traditional feature attribution for trees, such as split-based or gain-based measures, is inconsistent, meaning it can lower a feature’s assigned importance when the true impact of that feature increases.

Therefore, for the interpretation of the feature importance, we compute the SHAP (SHapley Additive exPlanation) values [21, 20]. SHAP is a widely used methodology. It is based on game theory [45], and local explanations [46], and it offers a means to estimate the contribution of each feature. Specifically, by combining many local explanations, models’ global behaviour can be structured, while retaining local faithfulness [47] to the original model, which generates detailed and accurate representations of the model’s behavior. It is important to underline that the relationship between the features and the target variable does not need to be causal, as SHAP could fail to answer causal questions accurately.

#### Explaining the global feature importance

In this study, we use SHAP as a tool to identify which features are the most important drivers of risk of inadequate micronutrient intake predictions, and how features drive the predictions. Considering that predictions are originally obtained at a household level, global feature importance presents the SHAP values across all households. This method computes the feature importance ranking for each trained model, explaining which features are the most important and the direction positive or negative, that is micronutrient inadequacy or adequacy, they drive the final prediction of each household. Overall, SHAP method significantly enhances our ability to generate explainable predictions, supporting both debugging and robustness, while also delivering interpretable results for relevant stakeholders.

## Acknowledgements

We would like to thank Kyriacos Koupparis for the continuous support and guidance. We are grateful to Hiwot Tadesse, Ekene Onyeagba, Rihanna Ibrahim, and Ahmad Ghaith for the support and feedback. We also thank all MIMI team members for the continuous feedback.

## Declarations

- Funding: This work was supported by the Gates Foundation (INV-037325) through the Modelling & mapping the risk of Inadequate Micronutrient Intake (MIMI) project.
- Conflict of interest/Competing interests: The authors have no conflicts of interest to declare
- Data availability: The data of the project are available upon request to the authors.
- Code availability: The code developed is available upon request to the authors.
- Author contribution: F.K. conceived the research, D.P. supervised the research, V.V. and D.P. designed the study, V.V. conducted the machine learning-related calculations, K.T. conducted the target-related calculations, V.V., K.T., I.L., M.J., G.M., S.D.P., F.K., D.P. interpreted and evaluated the findings. V.V., K.T., I.L., M.J., G.M., S.D.P., F.K., D.P. wrote and reviewed the manuscript.

## Appendix A

SHAP plots of the individual micronutrient models

Figures 5, 6, 7, 8, 9 provide the SHAP values of the models trained to predict the risk of each individual micronutrient for each country. Similar to the model trained to predict the overall risk (see Figure 2 in Section 4.2), we observe food group diversity-, education-, and climate-related features to be the most important. While the order of importance may vary between countries, the key features and their directional relationships remain largely consistent. As a general pattern, the plots reveal that for each specific micronutrient model the food group diversity features correspond to food items rich in that micronutrient. For instance, for models trained to predict the risk of inadequate folate intake, fruits emerge as the most important food group diversity feature in both countries (see Figure 6). For models trained to predict risk of inadequate vitamin B12 intake, meat, fish, and eggs are identified as the most important food group diversity features for both countries (see Figure 9). Overall, the results presented in the SHAP plots enhance model explainability and robustness.

**Figure 5:**
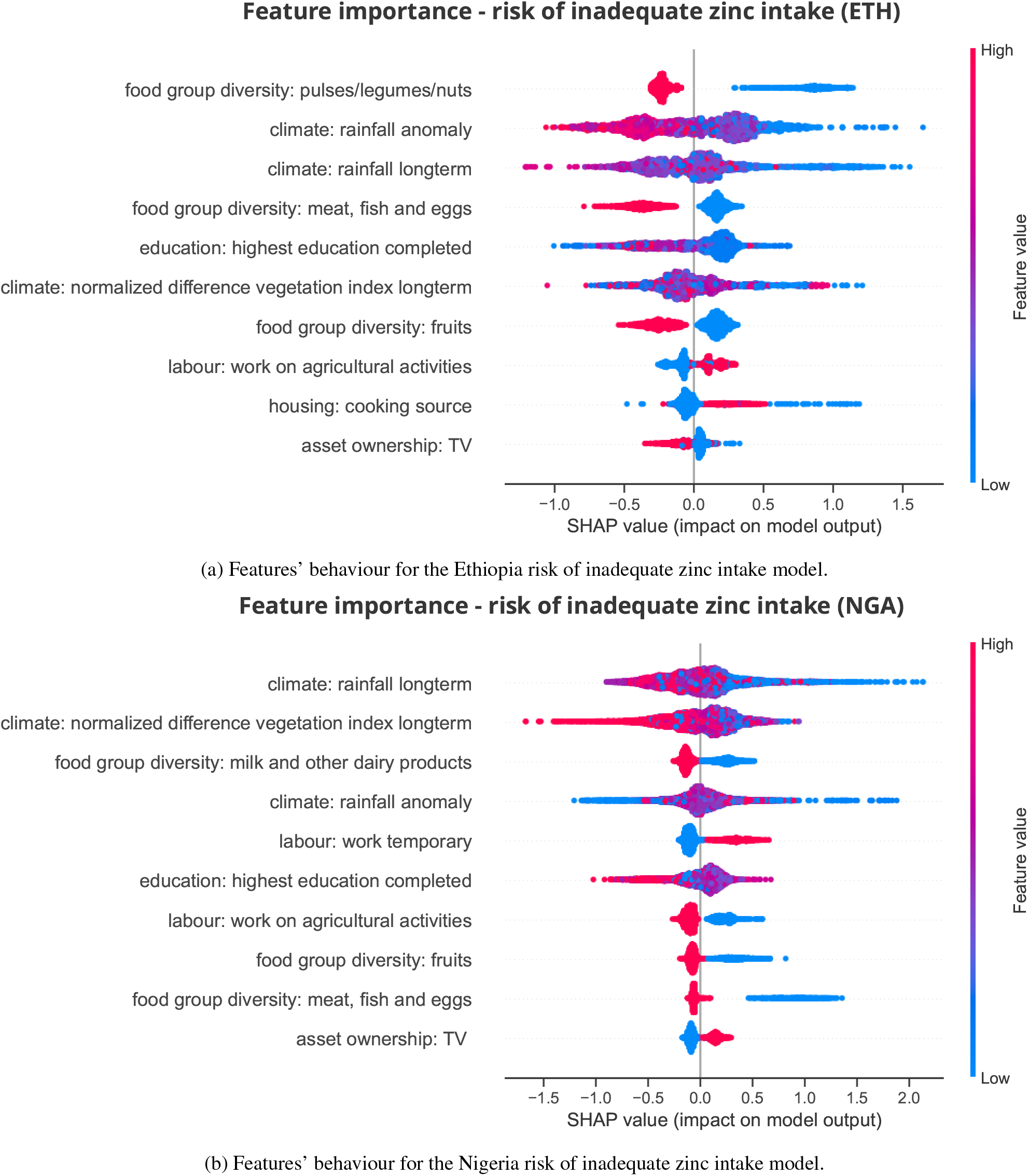
Features’ behaviour for the Ethiopia and Nigeria risk of inadequate zinc intake model.

**Figure 6:**
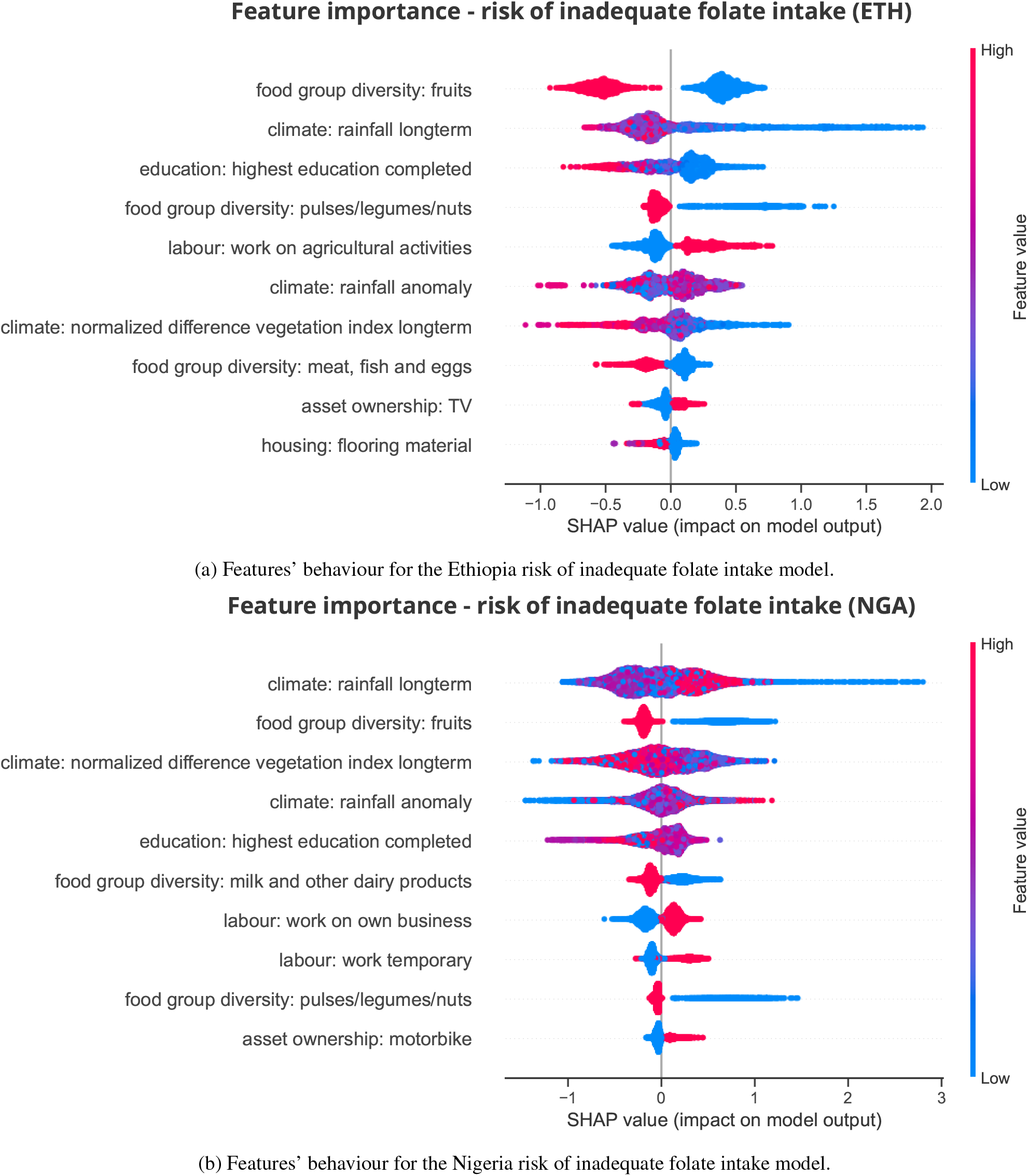
Features’ behaviour for the Ethiopia and Nigeria risk of inadequate folate intake model.

**Figure 7:**
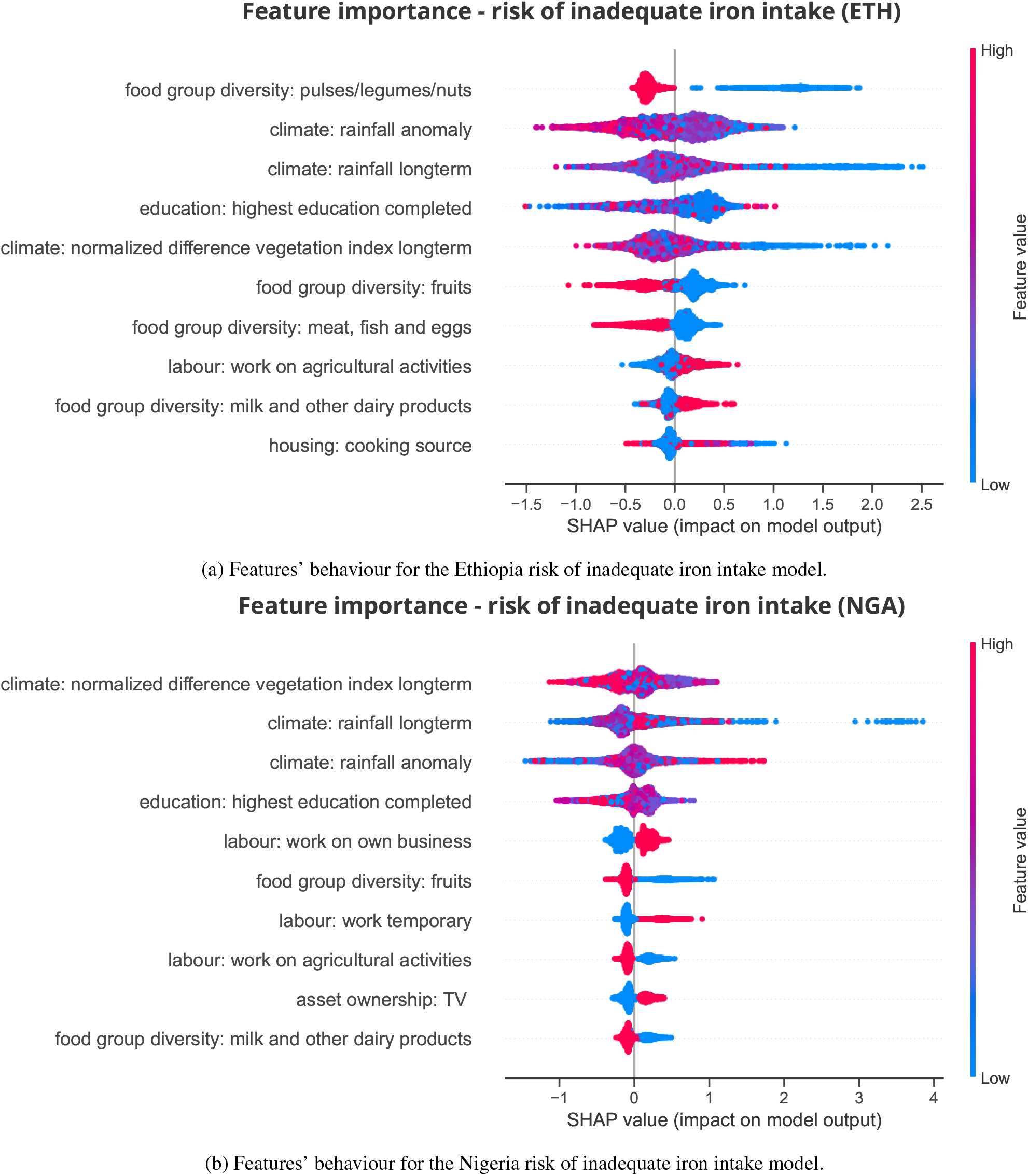
Features’ behaviour for the Ethiopia and Nigeria risk of inadequate iron intake model.

**Figure 8:**
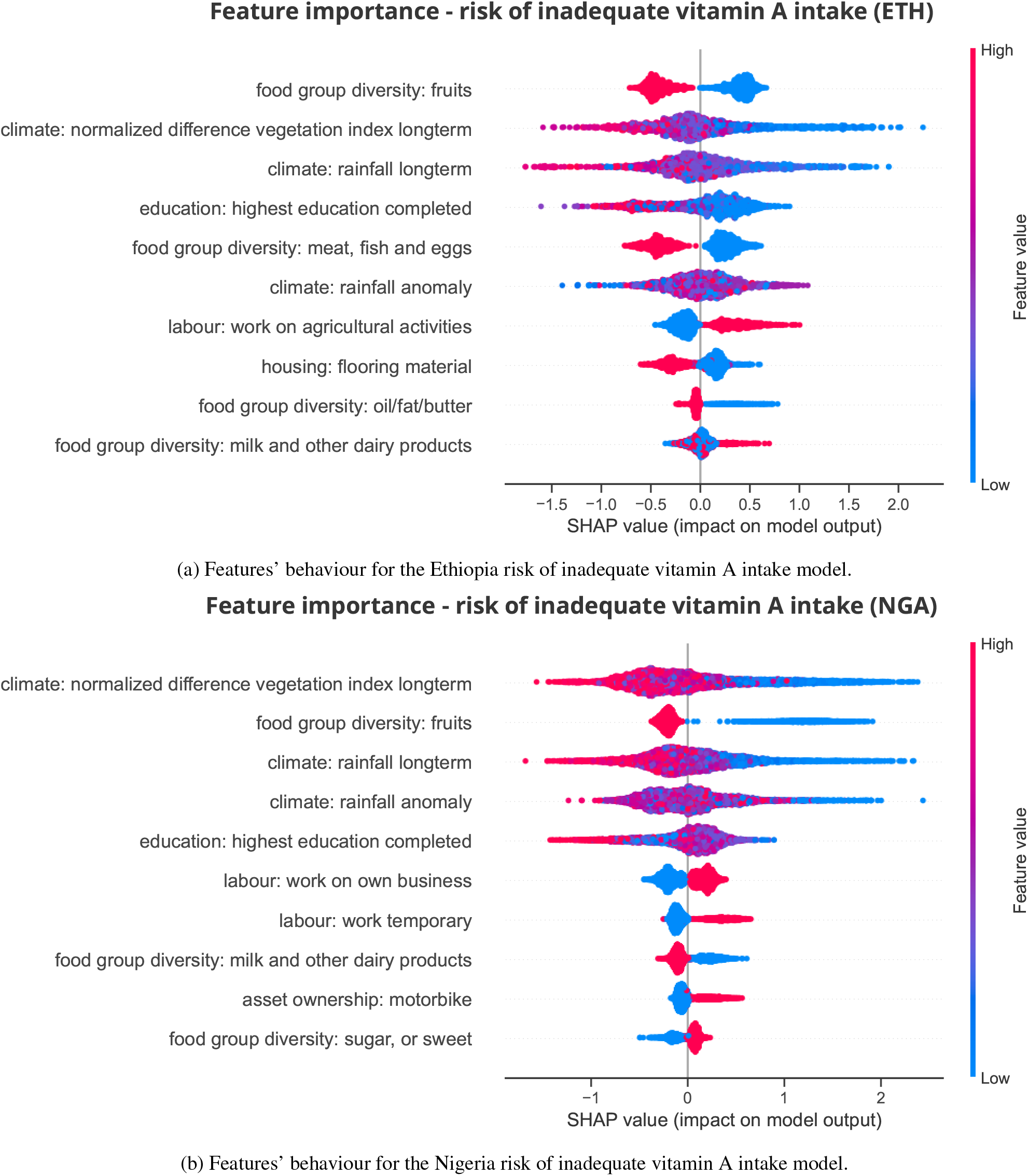
Features’ behaviour for the Ethiopia and Nigeria risk of inadequate vitamin A intake model.

**Figure 9:**
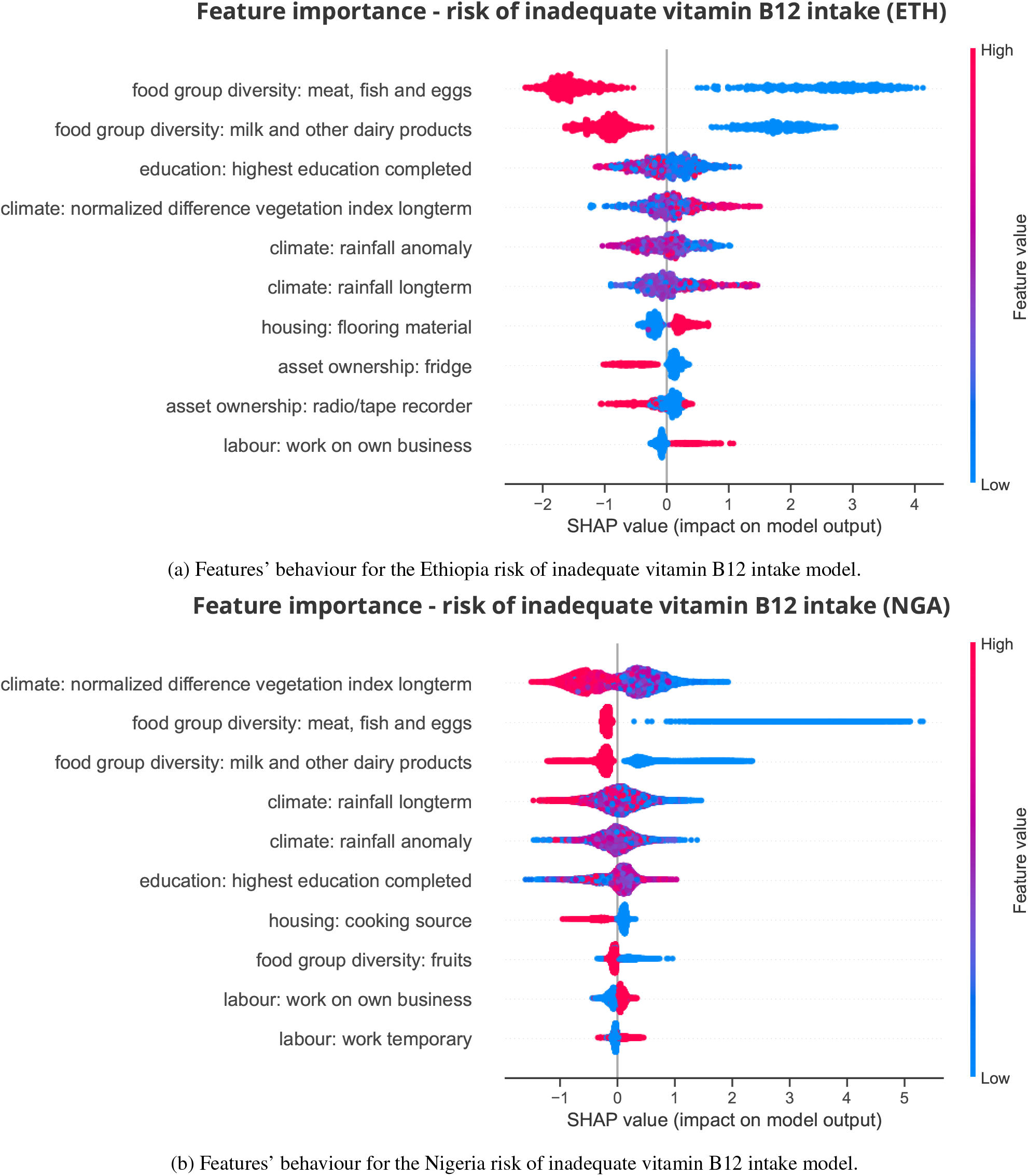
Features’ behaviour for the Ethiopia and Nigeria risk of inadequate vitamin B12 intake model.

## Appendix B

Selection of micronutrients and MAR

We focus on five micronutrients of particular public health importance, i.e., zinc, folate, iron, vitamin A, and vitamin B12 [2]. These micronutrients are selected based on multiple criteria: a) deficiencies of these micronutrients are commonly identified as being prevalent, and b) linked to dietary inadequacy (among other risk factors), with c) established links to critical health outcomes and d) they are often added to fortified cereal and staple grains for the first two reasons and as it is technically viable to do so [2]. First, we establish the scope of potential micronutrients based on relevance to large-scale fortification efforts, particularly those targeting staple foods and condiments distributed in social assistance programs. A shortlist of micronutrients is drawn from those included in international recommendations for wheat flour fortification [5] and specifications for rice fortification [48]. Second, micronutrients are selected based on their direct associations with critical health outcomes. Vitamin A and zinc are included due to their deficiencies being directly linked to increased child mortality through pathway of weakened immune function and heightened vulnerability to infectious diseases [49, 50]. Iron, folate, and vitamin B12 are included due to their deficiencies being major contributors to anaemia, which increases the risk of maternal mortality, complications during childbirth, and adverse pregnancy outcomes, including preterm birth and low birth weight (REF Fishman 2007). MAR is a practical indicator for programmes that address inadequate intake of multiple micronutrients simultaneously, such as large-scale food fortification. It also provides information on how adequately a diet meets essential nutrient requirements, helping assess the impact of nutrition programmes. MAR enables better comparisons across diverse populations and interventions, and helps policymakers and stakeholders to make informed decisions.

## Appendix C

WFP food groups for the dietary diversity score

As discussed in Section 4.2, we generate nine binary features from each Living Standards Measurement Study 2018/19 (LSMS). We align the food groups of the corresponding survey based on the Food Consumption Groups (FCGs), as defined by the United Nations World Food Programme (WFP) [39]. Table 3 presents the WFP FCGs, and examples of food groups from the Ethiopian and Nigerian LSMS surveys that fall under the FCGs.

## Appendix D

Matching of survey questions per category

We have worked on the consistency of the features by matching the same or similar questions from each country survey, for the purposes of feature engineering. Table 4 presents the socioeconomic categories, and the corresponding matched survey questions from the Living Standards Measurement Study 2018/19 (LSMS) both for Ethiopia (ETH) and Nigeria (NGA).

**Table 4:**
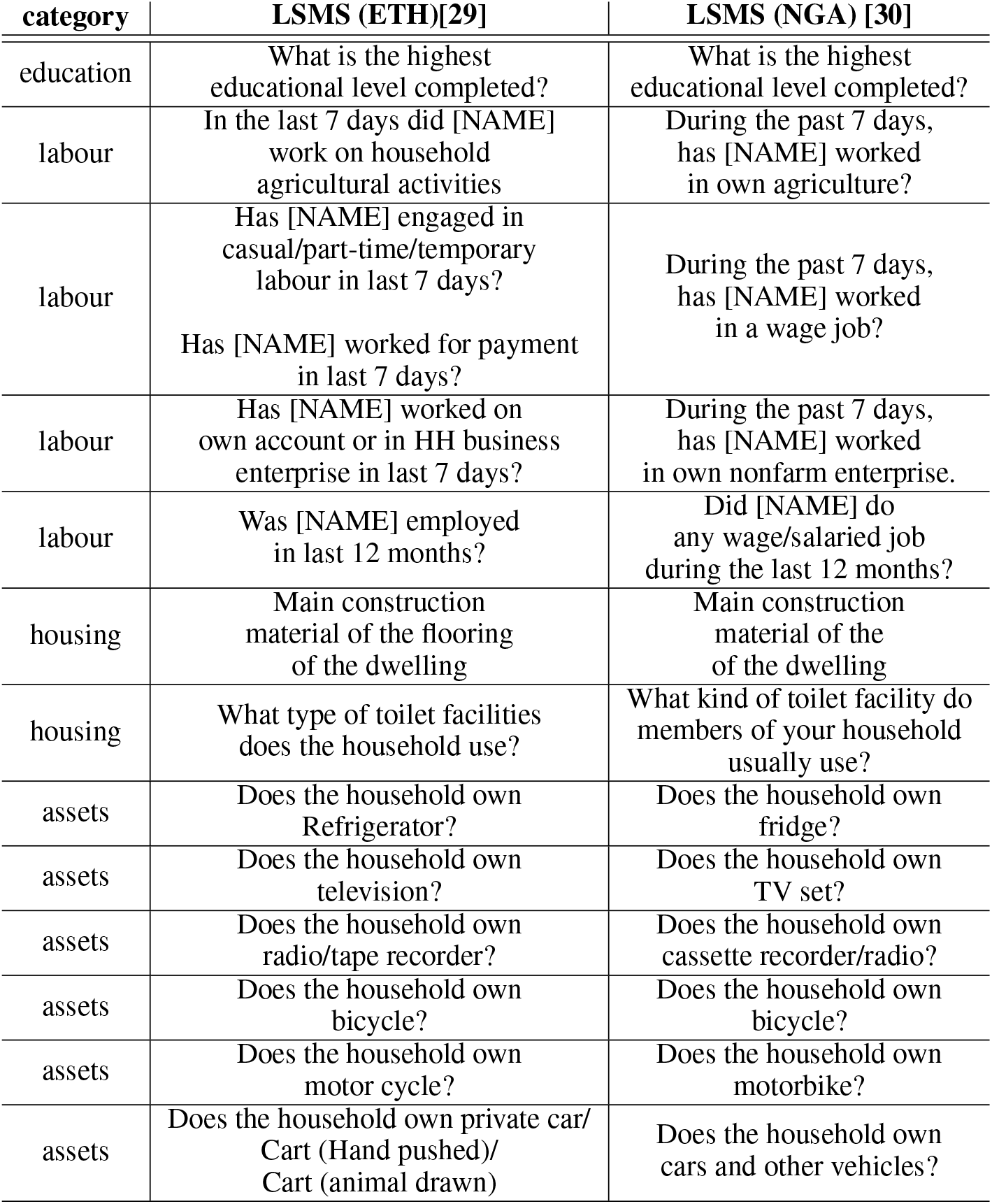
The table presents the socioeconomic categories, and the corresponding matched survey questions from the Living Standards Measurement Study 2018/19 both for Ethiopia (ETH) and Nigeria (NGA).

| category | LSMS (ETH)[29] | LSMS (NGA) [30] |
| --- | --- | --- |
| education | What is the highest educational level completed? | What is the highest educational level completed? |
| labour | In the last 7 days did [NAME] work on household agricultural activities | During the past 7 days, has [NAME] worked in own agriculture? |
| labour | Has [NAME] engaged in casual/part-time/temporary labour in last 7 days?<br>Has [NAME] worked for payment in last 7 days? | During the past 7 days, has [NAME] worked in a wage job? |
| labour | Has [NAME] worked on own account or in HH business enterprise in last 7 days? | During the past 7 days, has [NAME] worked in own nonfarm enterprise. |
| labour | Was [NAME] employed in last 12 months? | Did [NAME] do any wage/salaried job during the last 12 months? |
| housing | Main construction material of the flooring of the dwelling | Main construction material of the of the dwelling |
| housing | What type of toilet facilities does the household use? | What kind of toilet facility do members of your household usually use? |
| assets | Does the household own Refrigerator? | Does the household own fridge? |
| assets | Does the household own television? | Does the household own TV set? |
| assets | Does the household own radio/tape recorder? | Does the household own cassette recorder/radio? |
| assets | Does the household own bicycle? | Does the household own bicycle? |
| assets | Does the household own motor cycle? | Does the household own motorbike? |
| assets | Does the household own private car/ Cart (Hand pushed)/ Cart (animal drawn) | Does the household own cars and other vehicles? |

**Table 5:** Models’ hyperparameters and explored values. The hyper-parameters listed are tuned through 5-fold cross validation, exploring all possible combinations of the listed values.

| Hyperparameters | Values |
| --- | --- |
| *n estimators* | 100, 200, 500 |
| *learning rate* | 0.01, 0.05, 0.1 |
| *booster* | *gbtree, gblinear* |
| *gamma* | 0, 0.5, 1 |
| *alpha* | 0, 0.5, 1 |
| *lambda* | 0.5, 1, 5 |
| *maxdepth* | 3, 4, 5, 6 |

**Table 6:** Summary of the variables, the data sources, availability of data based on the country context.

| variables | data sources | availability |
| --- | --- | --- |
| targets | HCES data | data-rich contexts |
| food group diversity features | HCES data | data-rich contexts |
|  | WFP or other survey data | data-constrained contexts |
| socioeconomic features | HCES data | data-rich contexts |
|  | WFP or other survey data | data-constrained contexts |
| climate features | WFP’s Seasonal Explorer platform | both data-rich and data-constrained contexts |

## Appendix E

Distributions of the target variables

Figures 11-13 provide the distribution of all six target variables both for Ethiopia and Nigeria.

**Figure 10:**
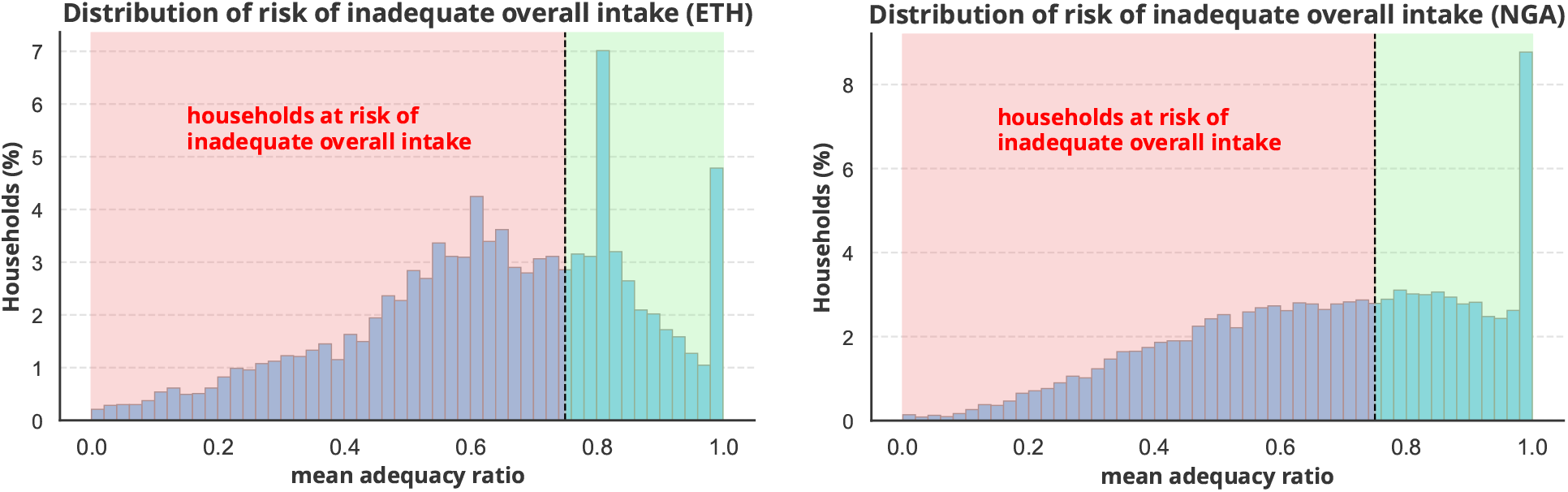
Distributions of the values of risk of inadequate overall intake, across households.

**Figure 11:**
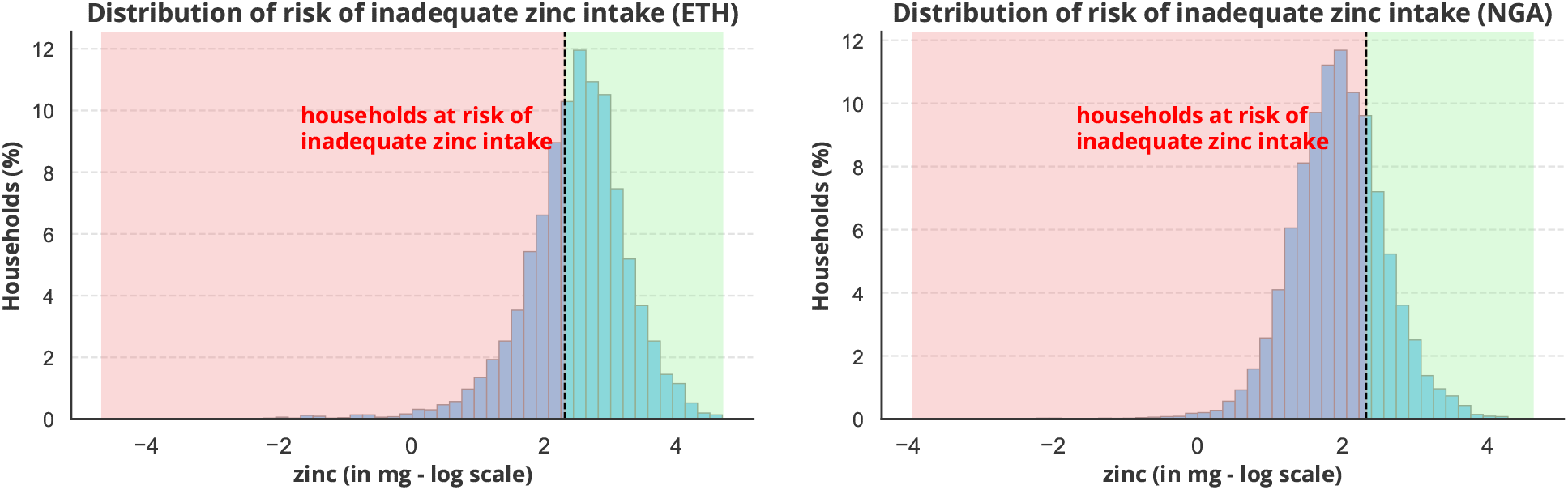
Distributions of the values of risk of inadequate zinc intake, across households.

**Figure 12:**
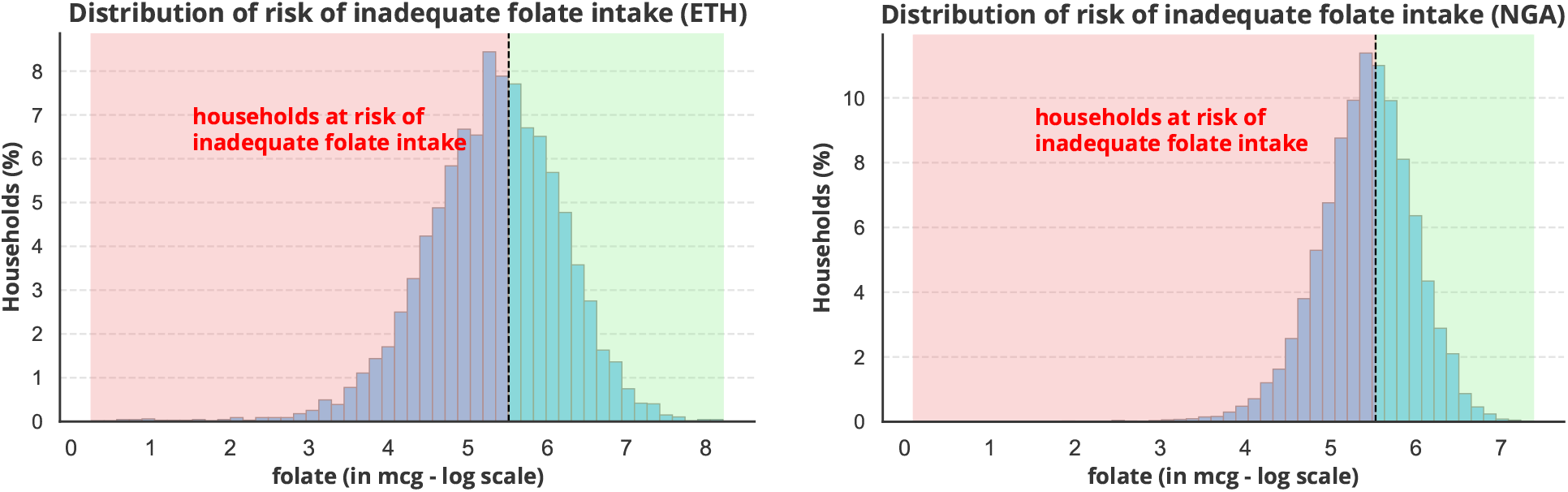
Distributions of the values of risk of inadequate folate intake, across households.

## Appendix F

Overestimations and underestimations of the models

As discussed in Section 2.3, due to differences in the distribution of some features the model trained in Nigeria and applied in Ethiopia overestimates the risk, whereas the model trained in Ethiopia and applied in Nigeria underestimates the risk. This can be explained by the feature importance and the SHAP values presented in Section 4.2. For example, for Ethiopia high values of feature “ndvi longterm” push towards adequacy, and medium/low values push towards inadequacy, although we notice some outliers. In Nigeria, low values of “ndvi longterm” push towards inadequacy, but high/medium values push both towards adequacy and inadequacy. Thus, the model trained in Ethiopia and applied in Nigeria underestimates the risk, considering there are some high/medium values that would push towards inadequacy. In contrast, the model trained in Ethiopia and applied in Nigeria underestimates the risk, considering that high/medium “ndvi longterm” values push mostly towards adequacy.

**Figure 13:**
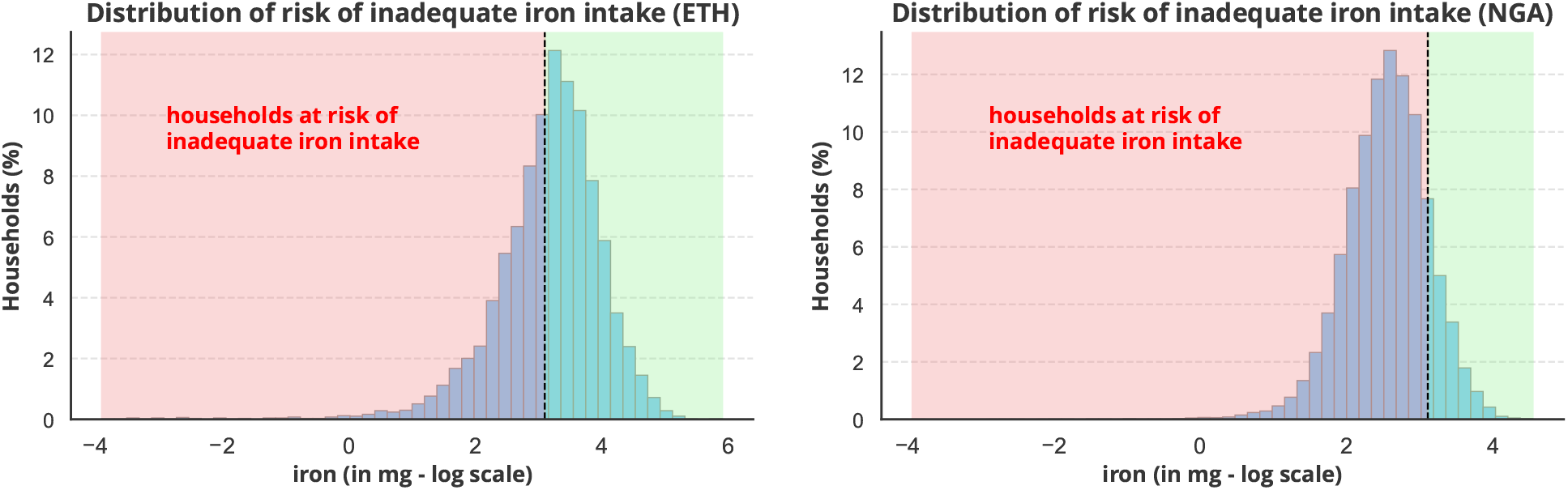
Distributions of the values of inadequate iron intake, across households.

**Figure 14:**
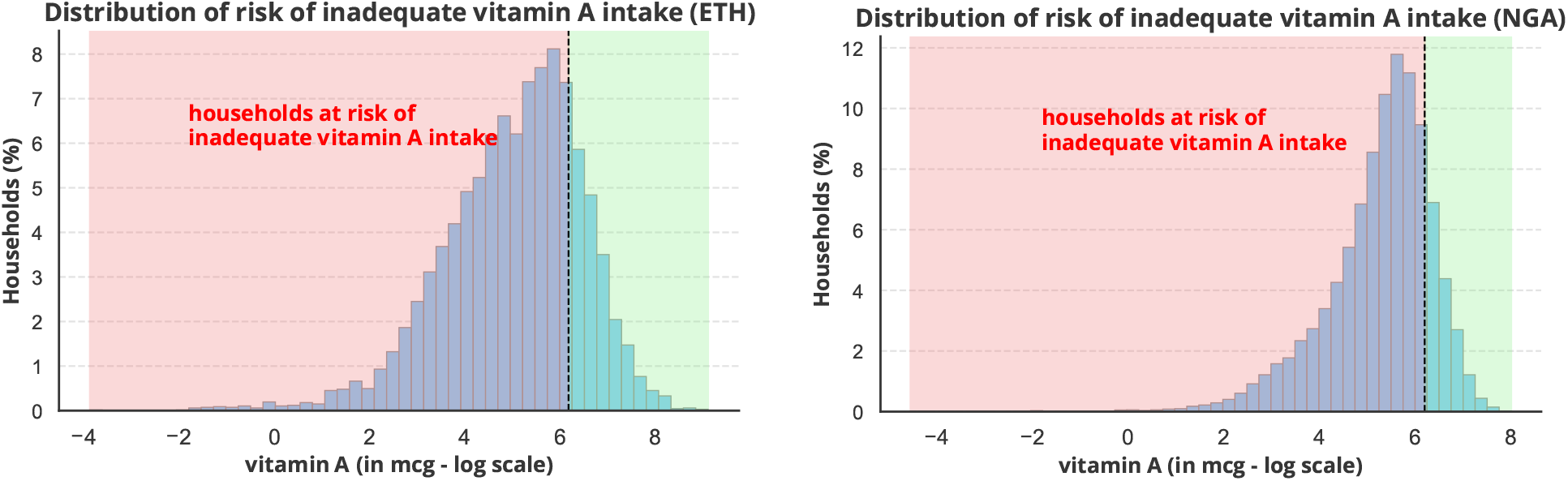
Distributions of the values of risk of inadequate vitamin A intake, across households.

**Figure 15:**
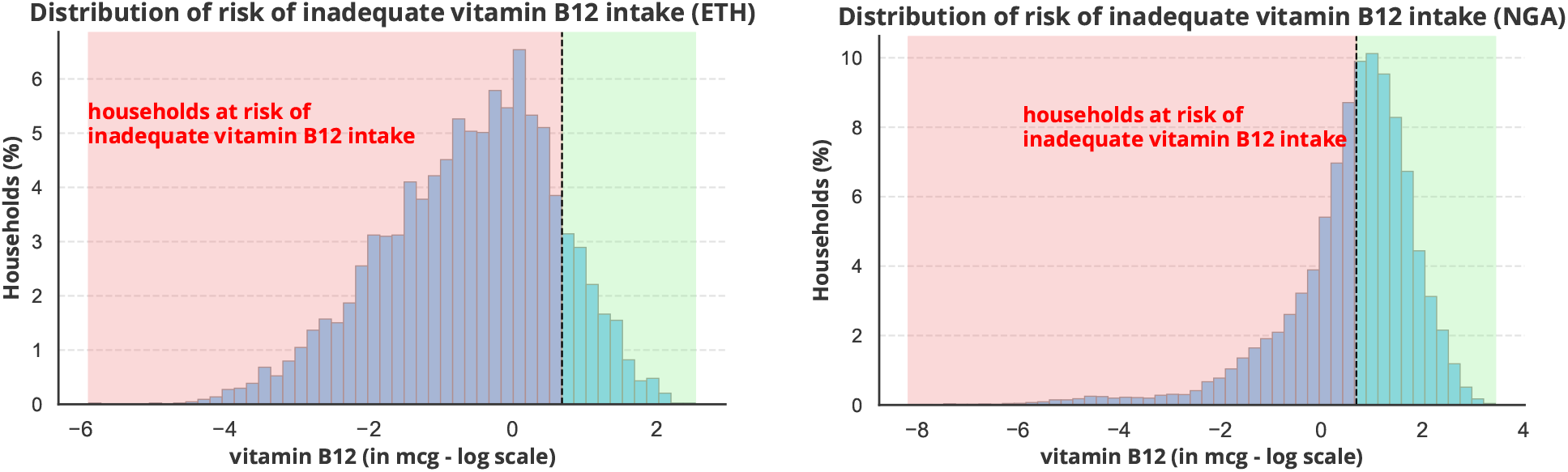
Distributions of the values of risk of inadequate vitamin B12 intake, across households.

## Appendix G

Scatter plots of the actual and predicted percentages of the risk of inadequate overall intake

Figure 16 presents the scatter plots of the actual and predicted percentages of risk of inadequate overall intake for the model trained in Ethiopia and applied in Nigeria (left) and for the model trained in Nigeria and applied in Ethiopia (right). Household level data is aggregated at first-level administrative unit resolution. Overall, we observe that the actual percentages are very similar to the predicted percentages, generated from the corresponding optimal probability thresholds (see Section 2.3), although for some first-level administrative units the models overestimate or underestimate the risk.

**Figure 16:**
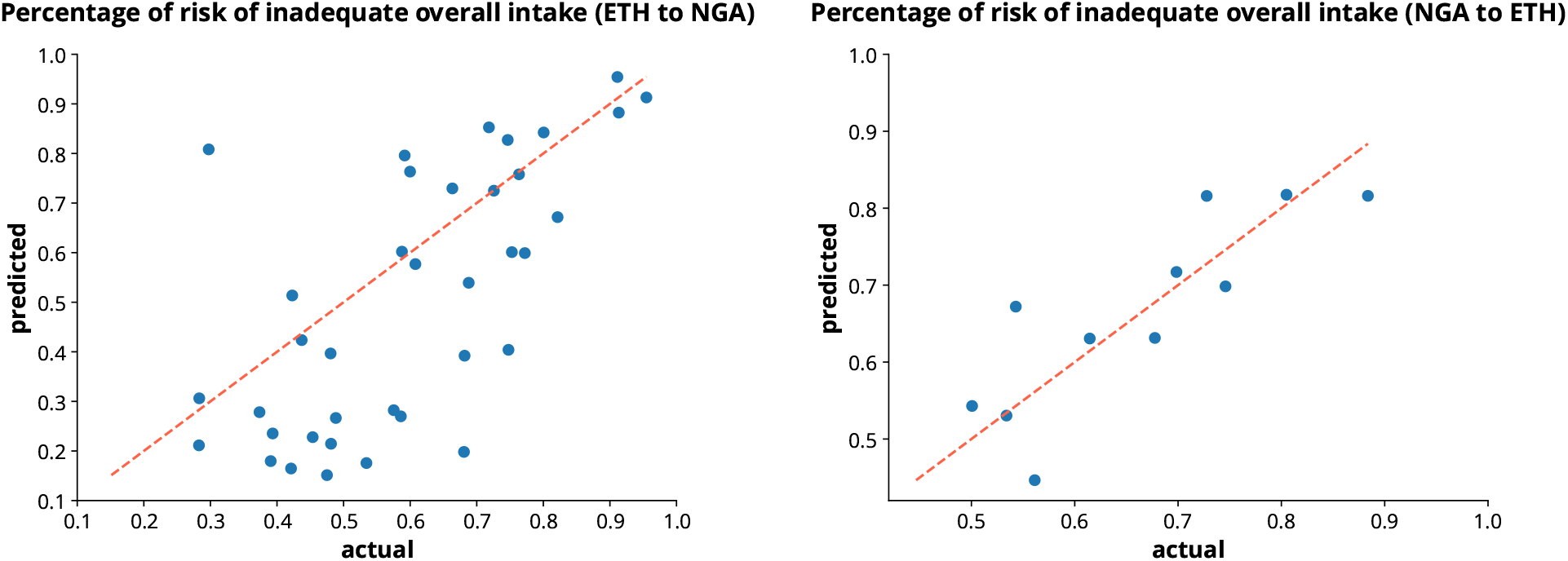
Scatter plots of the actual and predicted percentages of risk of inadequate overall intake for the model trained in Ethiopia and applied in Nigeria (left) and for the model trained in Nigeria and applied in Ethiopia (right). Data is aggregated at first-level administrative unit resolution.

## Appendix H

Extreme Gradient Boosting (XGBoost)

XGBoost [43] is a scalable machine learning classification system for tree boosting. It uses a gradient descent algorithm and incorporates a regularized model to prevent overfitting. XGBoost uses boosting, combining weak learners (usually decision trees with only one split, called decision stumps) sequentially, so that each new tree corrects the errors of the previous one. In particular, XGBoost corrects the previous mistakes made by the model, learns from it and its next step enhances the performance until there is no scope of further improvements. Its main advantage is that it is fast to execute and gives high accuracy.

## Appendix I

Hyperparameters

For XGBoost, we tune the *n estimators*, which accounts for the number of trees in the model, and the *maxdepth*, which is the maximum depth of the tree. We tune the *learning rate*, a value that in each boosting step, shrinks the weight of new features, preventing overfitting. Also, we tune the *booster*, which specifies the type of base learner used, such as *gbtree* for tree-based models or *gblinear* for linear models. We tune the *gamma*, that is the minimum loss reduction required to make a further partition on a leaf node of the tree. Last, we tune *alpha* and *lambda*, which are the *L*1 and *L*2 regularization terms on weights, respectively.

## Appendix J

Wealth, rural-urban, and geographic stratifications

For the purposes of this study, households are stratified by wealth quintile, rural-urban area, and geography. Wealth is defined as quintiles of inflation-adjusted total per capita household expenditure, derived from the corresponding Household Consumption and Expenditure Surveys (HCES) collected as a part of the Living Standards Measurement Studies 2018/19. Similarly, households are stratified as rural or urban, based on the corresponding HCES data. Also, households are stratified by geography (i.e., regions) at first-level administrative unit resolution. Actual and predicted risk (and/or absence of risk) of inadequate micronutrient intake, is then aggregated by wealth quintile, by rural-urban area, and by geography. Actual and predicted data are weighted with the calculated weights for maintaining representativeness (see details in K).

## Appendix K

Survey weights

For wealth quintile, rural-urban, and geographic stratifications, and visualisations we use the weights provided from the corresponding Household Consumption and Expenditure Surveys (HCES) collected as a part of the Living Standards Measurement Studies 2018/19. The weights are calculated to maintain representativeness over the national-level population of rural-urban areas, as well as the regional population. They reflect the adjusted probability of selecting the household in the sample. We apply the weights at a household level which sum up to the population of households. More details on the methodologies followed can be found in the data documentation of the studies [29, 30].

## Appendix L

Variables, data sources and availability in each country context

Table 6 presents the variables, indicative data sources and availability in each country context. In data-constrained country contexts Household Consumption and Expenditure Surveys (HCES) data are not available and thus target data cannot be estimated. In such contexts, other available survey data can be used to generate food group diversity, and socioeconomic features, while climate features can be still extracted from the WFP’s Seasonal Explorer platform [27].

## Footnotes

1 Fortification is the practice of deliberately increasing the content of one or more micronutrients (i.e., vitamins and minerals) in a food or condiment to improve the nutritional quality of the food supply and provide a public health benefit with minimal risk to health [5].

